# Inflammation promotes adipocyte lipolysis via IRE1 kinase

**DOI:** 10.1101/2020.04.07.030148

**Authors:** Kevin P. Foley, Yong Chen, Nicole G. Barra, Mark Heal, Kieran Kwok, Akhilesh K. Tamrakar, Wendy Chi, Brittany M. Duggan, Brandyn D. Henriksbo, Yong Liu, Jonathan D. Schertzer

## Abstract

Obesity associates with inflammation, insulin resistance and higher blood lipids. It is unclear if immune responses facilitate lipolysis separate from hormone or adrenergic signals. We found that an ancient component of ER stress, inositol-requiring protein 1 (IRE1), discriminates inflammation-induced adipocyte lipolysis versus lipolysis regulated by adrenergic or hormonal stimuli. Inhibiting IRE1 kinase activity was sufficient to block adipocyte-autonomous lipolysis from multiple inflammatory ligands, including bacterial components, certain cytokines, and thapsigargin-induced ER stress. Adipocyte-specific deletion of IRE1 in mice prevented inflammatory ligand-induced lipolysis in adipose tissue. IRE1 kinase activity was dispensable for isoproterenol and cAMP-induced lipolysis in adipocytes and mouse adipose tissue. IRE1 RNase activity was not associated with inflammation-induced adipocyte lipolysis. We found no role for canonical unfolded protein responses (UPR) or ABL kinases in linking ER stress to lipolysis. Lipolysis was unchanged in adipose tissue from GRP78/BiP^+/-^ compared to littermate mice. Tyrosine kinase inhibitors (TKIs) such as imatinib, which reduce ER stress and IRE1 RNase activity, did not alter lipolysis from inflammatory stimuli. Inhibiting IRE1 kinase activity blocked adipocyte NF-κB activation and Interleukin-6 (Il6) production due to inflammatory ligands. Inflammation-induced lipolysis mediated by IRE1 occurred independently from changes in insulin signalling in adipocytes. Therefore, inflammation can promote IRE1-mediated lipolysis independent of adipocyte insulin resistance. Our results show that IRE1 propagates an inflammation-specific lipolytic program independent from hormonal or adrenergic regulation, including insulin resistance. Targeting IRE1 kinase activity may benefit metabolic syndrome and inflammatory lipid disorders.

**Significance:** Adipocytes maintain metabolic homeostasis by storing nutrients and releasing lipids into the blood via lipolysis. Catecholamines stimulate adrenergic-mediated lipolysis, whereas insulin inhibits lipolysis. Obesity is associated with elevated blood lipids and inflammation, which can impair insulin-mediated suppression of lipolysis (i.e. insulin resistance). It is unclear if inflammatory triggers of lipolysis require insulin resistance or if specific lipolytic triggers engage distinct cell stress components. We found that a specific ER stress response was required for inflammation-mediated lipolysis, not adrenergic-mediated lipolysis. Bacterial and cytokine-induced lipolysis required adipocyte IRE1 kinase activity, but not IRE1 RNase activity typical of the ER stress-related unfolded protein response. We propose that inflammatory triggers of lipolysis engage IRE1 kinase independent of catecholamine and hormone responses, including insulin resistance.

**Graphical Abstract:** 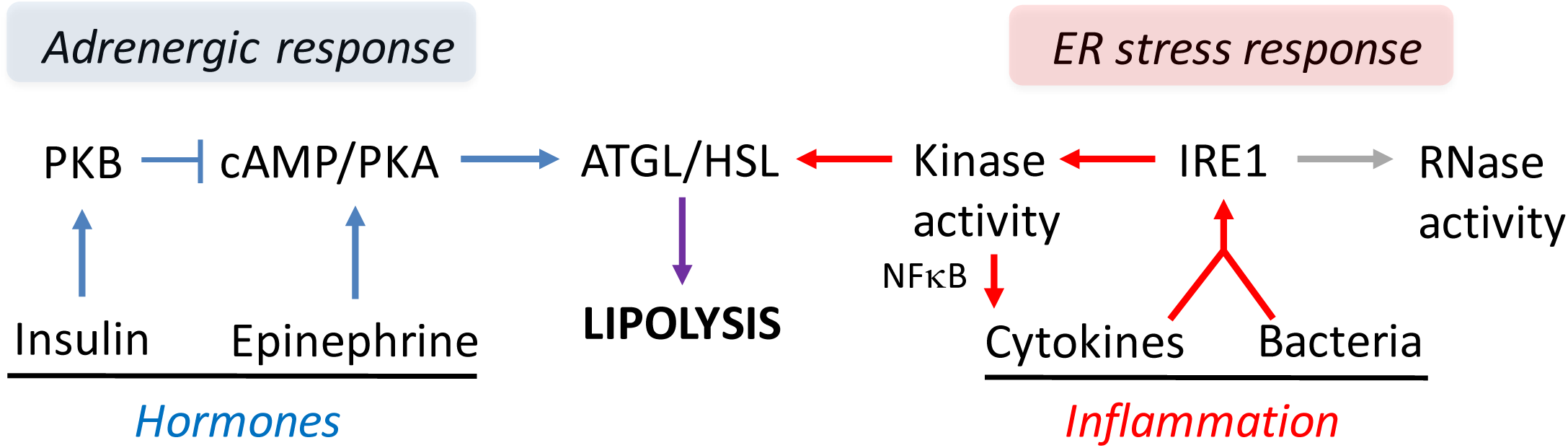

IRE1 kinase activity promotes an inflammation-specific adipocyte lipolytic program that is separate from hormonal or adrenergic regulation of lipolysis.

## Introduction

Obesity and metabolic diseases are pervasive (1, 2). Some consequences of obesity are underpinned by compartmentalized immune responses in tissues that participate in metabolic regulation. This “metaflammation” can manifest from autocrine responses within metabolic cell types that regulate glucose and lipid metabolism. Paracrine communication between immune and metabolic cells can also impair local and systemic endocrine and metabolic responses (3–5). For example, inflammation of metabolic tissues can manifest as insulin resistance (6). However, the cellular nodes that dictate metabolic defects linked to inflammation-specific responses are ill-defined.

Adipocyte expansion during obesity coincides with increased adipose tissue inflammation, including aberrant cytokine production from both adipocytes and adipose-resident immune cells. Obesity and inflammation in the adipose tissue environment are also associated with adipocyte insulin resistance, including reduced ability of insulin to inhibit lipolysis. There is a bi-directional link between lipolysis and adipocyte inflammation. Inflammatory mediators, such as tumor necrosis factor (TNF) and interleukin (Il)6 can increase adipocyte lipolysis. Elevated levels of saturated free fatty acids (7, 8) and even lipolysis itself can promote cellular and tissue inflammation (9).

Excessive lipolysis can result in the accumulation of ectopic lipids in the liver and skeletal muscles, which can subsequently contribute to systemic insulin resistance (10). Increased lipolysis during obesity is often formulated in terms of adipocyte insulin resistance. However, inflammation-induced lipolysis can occur independently of changes in insulin action (11–13). More importantly, it is unclear if certain cellular triggers are specific to inflammation-induced lipolysis compared to adrenergic control of lipolysis, which governs adipocyte metabolism and lipid flux via catecholamines and other hormones. We hypothesized that only inflammation engaged ancient cellular stress responses to provoke lipolysis, independently of hormonal influence on lipolysis, including insulin resistance.

Seminal work discovered how components of the endoplasmic reticulum (ER) stress response link inflammation to defects in glucose and lipid metabolism (14, 15). For example, nitrosylation of inositol-requiring protein 1 (IRE1) can compromise the unfolded protein response (UPR) and promote insulin resistance in obese mice (16). ER stress components include IRE1, activating transcription factor 6 (ATF6) and protein kinase RNA-like ER kinase (PERK). Much of the work on metabolic disease and ER stress has focussed on the UPR aspects of these components. For example, IRE1 can integrate stress signals from changes in cellular metabolism and nutrient levels coincident with overload of misfolded proteins (17). We questioned whether ER stress could propagate stress from inflammatory cues into a lipolytic program in adipocytes and whether the UPR or other kinases were involved. It was prudent to consider the kinase activity of ER stress components in the regulation of lipolysis because thapsigargin, tunicamycin, and brefeldin A, which promote ER stress through different pathways, all engage extracellular signal–regulated kinase (ERK)2 to promote hormone sensitive lipase (HSL)-mediated lipolysis (18). Inflammatory triggers of lipolysis may work in a similar ER-stress dependent manner, since bacterial lipopolysaccharide (LPS) and peptidoglycan (PGN) engage ERK2 and other kinases to promote lipolysis in adipocytes (11, 12). Recent evidence has also shown that ERK2 promotes serine 247 phosphorylation and activation of the Beta3 adrenoreceptor and increased adipocyte lipolysis during obesity (19). This seminal work showed that adrenergic stimuli engaged an ERK2-mediated stress response to promote lipolysis (19). Thus, kinases can communicate with adrenergic responses to promote lipolysis. However, it is not clear if inflammatory triggers of lipolysis, such as bacteria or cytokines, require a specific kinase to promote lipolysis. There are clearly links between ER stress, inflammation and increased lipolysis during obesity. We sought to define if a kinase inherent to ER stress discriminates immune versus hormonal regulation of lipolysis.

## Results

### IRE1 kinase activity mediates inflammation-induced lipolysis

To determine if ER stress propagated adipocyte lipolysis from an inflammatory stimulus, differentiated 3T3-L1 adipocytes were treated with or without 10 µg/mL NOD1 ligand (FK565) and increasing concentrations of the chemical chaperons TUDCA or 4-PBA (Figure 1A, B). FK565, a synthetic mimetic of bacterial cell wall peptidoglycan (PGN) typical of gram-negative bacteria, stimulated a ∼5-fold increase in glycerol release rate which was dose-dependently inhibited by TUDCA and 4-PBA (Figure 1A-B). These results indicate that some aspect of ER stress participates in bacterial cell wall stimulated adipocyte lipolysis. We next examined if ER stress was required for induction of lipolysis with multiple inflammatory ligands compared to lipolysis caused by the adrenergic stimulus, isoproterenol. We used the inflammatory stimuli PGN (FK565 10 µg/L = NOD1 activation), LPS (500 ng/mL = TLR4 activation), TNF (10 ng/mL = paracrine cytokine), or thapsigargin (1 µM = ER stress activation), all of which increased the glycerol release rate in 3T3-L1 adipocytes. The increased lipolysis from all of these stimuli was blocked by 1 mg/mL TUDCA (Figure 1C). However, glycerol release induced by the β-adrenergic agonist isoproterenol (2 µM) was not altered by TUDCA (Figure 1C). These results suggest that ER stress may discriminate lipolysis from inflammation compared to catecholamines. We next tested the sensitivity of FK565, thapsigargin, and isoproterenol stimulated lipolysis to ATGL inhibition using Atglistatin in 3T3-L1 adipocytes. Inhibition of ATGL with Atglistatin blocked lipolysis induced by each of these ligands, confirming that glycerol release marked canonical activation of an ATGL-sensitive lipolytic program in adipocytes (Figure 1D).

**Figure 1:**
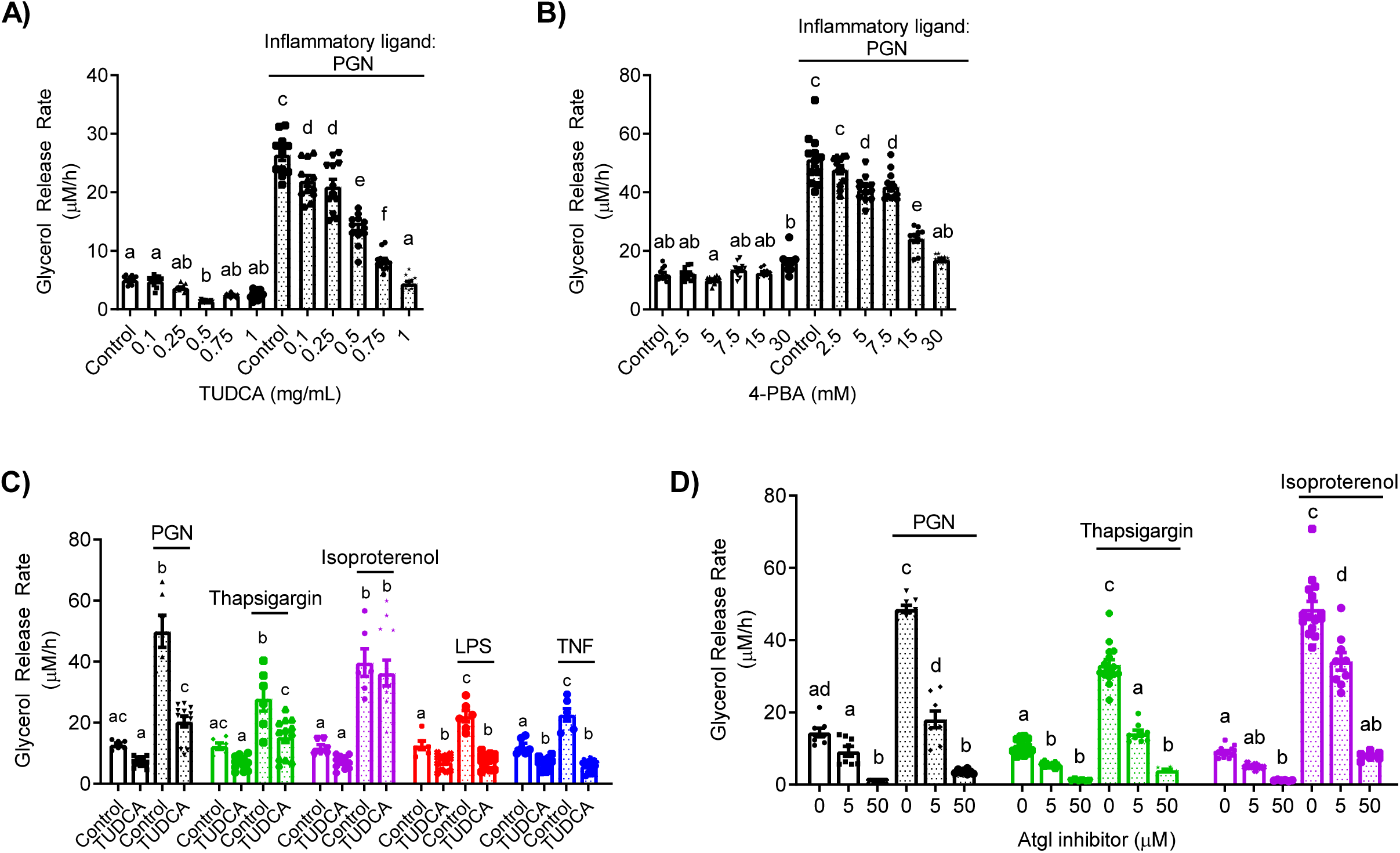
ER-stress mediates inflammation-induced lipolysis. A-B) Glycerol release rates (µM/h) in 3T3-L1 adipocytes treated with increasing concentrations of the chemical chaperones TUDCA (A; N=12) or 4-PBA (B; N=10-12) in the absence (white bars) or presence (dotted bars) of the inflammatory ligand peptidoglycan (PGN) using 10 µg/mL FK565 C) Glycerol release rates in 3T3-L1 adipocytes measured in the presence or absence of 1 mg/mL TUDCA, treated with vehicle (mock-treatment), 10 µg/mL PGN, 1 µM thapsigargin, 2 µM isoproterenol, 500 ng/mL LPS, or 10 ng/mL TNF (N=6-12). D) Glycerol release rates in 3T3-L1 adipocytes treated with increasing concentrations of the ATGL inhibitor Atglistatin in the absence or presence of 10 µg/mL PGN, 1 µM thapsigargin or 2 µM isoproterenol (N=6-15). Values are mean ± SEM. Statistical significance was measured as p<0.05 using two-way ANOVA. Post hoc analysis was performed using Tukey’s multiple comparisons test. Conditions with different letters (a, b) denote a statistical difference compared to all other conditions without the same letter.

IRE1 has been shown to link ER stress to NOD1/2-mediated immune responses (20, 21). We examined if IRE1 was the component of ER stress required for lipolysis stimulated by multiple inflammatory factors compared to adrenergic/hormonal pathways. Glycerol release was measured in 3T3-L1 cells treated with KirA6, an IRE1 kinase inhibitor (which also attenuates RNase activity) or one of two IRE1 RNase inhibitors, KirA8 and 4µ8C (Figure 2A-C). KirA6 dose dependently lowered PGN-mediated lipolysis, where 1 μM KirA6 blocked an increase in lipolysis (Figure 2A). KirA8 and 4µ8C did not lower glycerol release (Figure 2B, C). We also confirmed that 10 µM KirA8 blocked the RNase activity of IRE1 evinced by lower levels of spliced XBP1 in adipocytes treated with thapsigargin (Figure 2D). These data suggest IRE1 kinase activity (but not RNase activity) propagates inflammation-induced adipocyte lipolysis from a bacterial cell wall component. To test the role of IRE1 in mediating lipolysis induced by other inflammatory ligands, differentiated 3T3-L1 cells were treated with or without the IRE1 inhibitor KirA6 (1 µM) for 1 hour before treatment with the inflammatory stimuli PGN, LPS, TNF or thapsigargin. KirA6 completely blocked lipolysis stimulated by each of these inflammatory stimuli (Figure 2E). In contrast, KirA6 did not inhibit lipolysis stimulated by the hormonal stimulus isoproterenol (2 µM) (Figure 2E). We also confirmed that KirA6 did not change 8-Br-cAMP-induced lipolysis in adipocytes (Figure 2F). These data demonstrate that IRE1 kinase activity is required for inflammatory, but not adrenergic-cAMP-stimulated, lipolysis in adipocytes.

**Figure 2:**
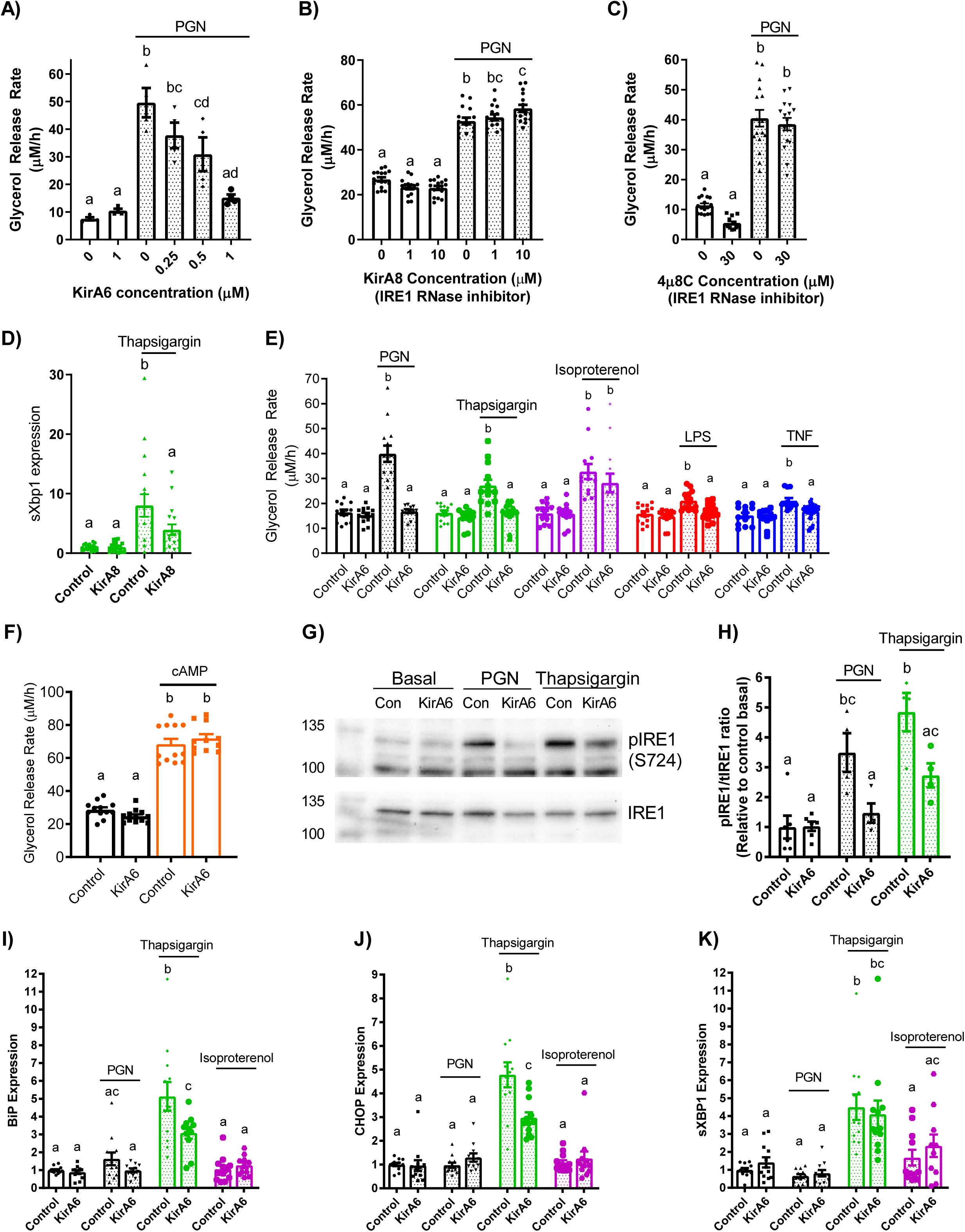
Inflammatory ligands stimulate lipolysis via IRE1 kinase not RNase activity. A-C) Glycerol release rates (µM/h) in 3T3-L1 adipocytes treated with increasing concentrations of the IRE1 inhibitor KirA6 (A; N=4), the IRE1 RNase inhibitor KirA8 (B; N=16), or the IRE1 RNase inhibitor 4µ8C (C; N=12-16). D) Expression of spliced Xbp1 in 3T3-L1 adipocytes treated with 1 μM Thapsigargin for 3 hours with and without 10 μM KirA8. E) Glycerol release rates in differentiated 3T3-L1 adipocytes treated with or without 1 µM KirA6 (IRE1 inhibitor) and one of vehicle (mock-treatment), 10 µg/mL FK565 (PGN), 1 µM thapsigargin (ER stress activator), 2 µM isoproterenol (β-adrenergic agonist), 500 ng/mL LPS (TLR4 activator), or 10 ng/mL TNF, as indicated (N=12-14). F) Glycerol release rates in 3T3-L1 adipocytes treated with 1 μM KirA6 and 0.5 mM 8-Br-cAMP (N=10-12). G) Phosphorylation of IRE1 was measured in 3T3-L1 adipocytes after stimulation with vehicle, PGN (10 µg/mL), or thapsigargin (1 µM) for 3 hours, with or without 1 µM KirA6 (N=4-6). Blots were stripped and re-probed for total IRE1. H) Quantification of Western blots for the ratio between phosphorylated IRE1 and total IRE1, expressed relative to the basal control sample. I-K) RNA isolated from 3T3-L1 adipocytes was treated for 3 hours with vehicle, FK565 (PGN, 10 µg/mL), thapsigargin (1 µM), or isoproterenol (2 µM) in the absence or presence of KirA6 (1 µM). Expression of BiP (I), CHOP (J), and spliced Xbp1(K) (N=12). Values are mean ± SEM. Statistical significance was measured as p<0.05 using two-way ANOVA. Post hoc analysis was performed using Tukey’s multiple comparisons test. Conditions with different letters (a, b) denote a statistical difference compared to all other conditions without the same letter.

We next examined IRE1 phosphorylation at Serine-724 (to biomark kinase activity) and transcript levels of key UPR intermediates (that are regulated by RNase activity). Treatment of 3T3-L1 adipocytes with PGN or thapsigargin for 3 hours increased levels of phosphorylated IRE1 (S724) relative to total levels of IRE1 (tIRE1) in 3T3 adipocytes, an effect that was inhibited by 1μM KirA6 (Figure 2G, H). After this same 3 hour treatment, thapsigargin, but not PGN or isoproterenol, increased expression of BiP, CHOP, and sXBP1 (Figure 2I-K). These data show that neither bacterial ligand (i.e. PGN) nor adrenergic stimulus (isoproterenol) increase ER stress markers typical of the UPR in adipocytes. Further, the increase in expression of BiP and CHOP induced by thapsigargin was sensitive to KirA6, whereas higher levels of spliced XBP1 (sXBP1), a key target of IRE1 RNase activity, were not affected by KirA6. These data support the concept that IRE1 kinase activity rather than RNase activity regulates lipolysis induced by inflammatory, but not hormonal, stimuli, independent of canonical UPR signalling in adipocytes.

We next measured glycerol release in mouse-derived epidydimal adipose tissue explants. Adipose tissue from wild type (WT) C57Bl6/J mice showed that PGN (10 µg/mL) and isoproterenol (2 µM) increased glycerol release, but only the increased lipolysis caused by PGN was inhibited by KirA6 (Figure 3A, B). KirA6 did not alter isoproterenol stimulated lipolysis in adipose tissue (Figure 3B). We then tested if the canonical UPR was involved in ER stress or IRE1-related lipolysis using Grp78^+/-^ and WT littermate mice. We found that neither TUDCA nor KirA6 altered glycerol release in adipose tissue explants derived from Grp78^+/-^ mice, which had comparable lipolysis to WT mice (Figure 3C). These data support our observations in 3T3-L1 adipocytes that IRE1 regulates lipolysis induced by inflammation independent of canonical UPR signalling, and that IRE1 does not alter lipolysis from adrenergic stimuli in adipose tissue. Finally, we generated mice with adipocyte-specific knockout of IRE1 (*Ern1* gene), where protein levels of IRE1 were lower/absent in adipocytes from epidydimal white adipose tissue (epiWAT), but not in the stromal vascular fraction (SVF), liver or muscle from these mice (Figure 3D). Adipose tissue explants from these IRE1 adipocyte-specific knockout mice (AKO) had lower glycerol release in response to thapsigargin or LPS when compared with Floxed control littermate mice (Figure 3E, F). This data reinforces our findings using chemical inhibitors of IRE1 (i.e. KirA6) in 3T3-L1 adipocytes and provides direct genetic evidence that IRE1 mediates lipolysis induced by inflammatory or bacterial ligands in mouse adipose tissue.

**Figure 3:**
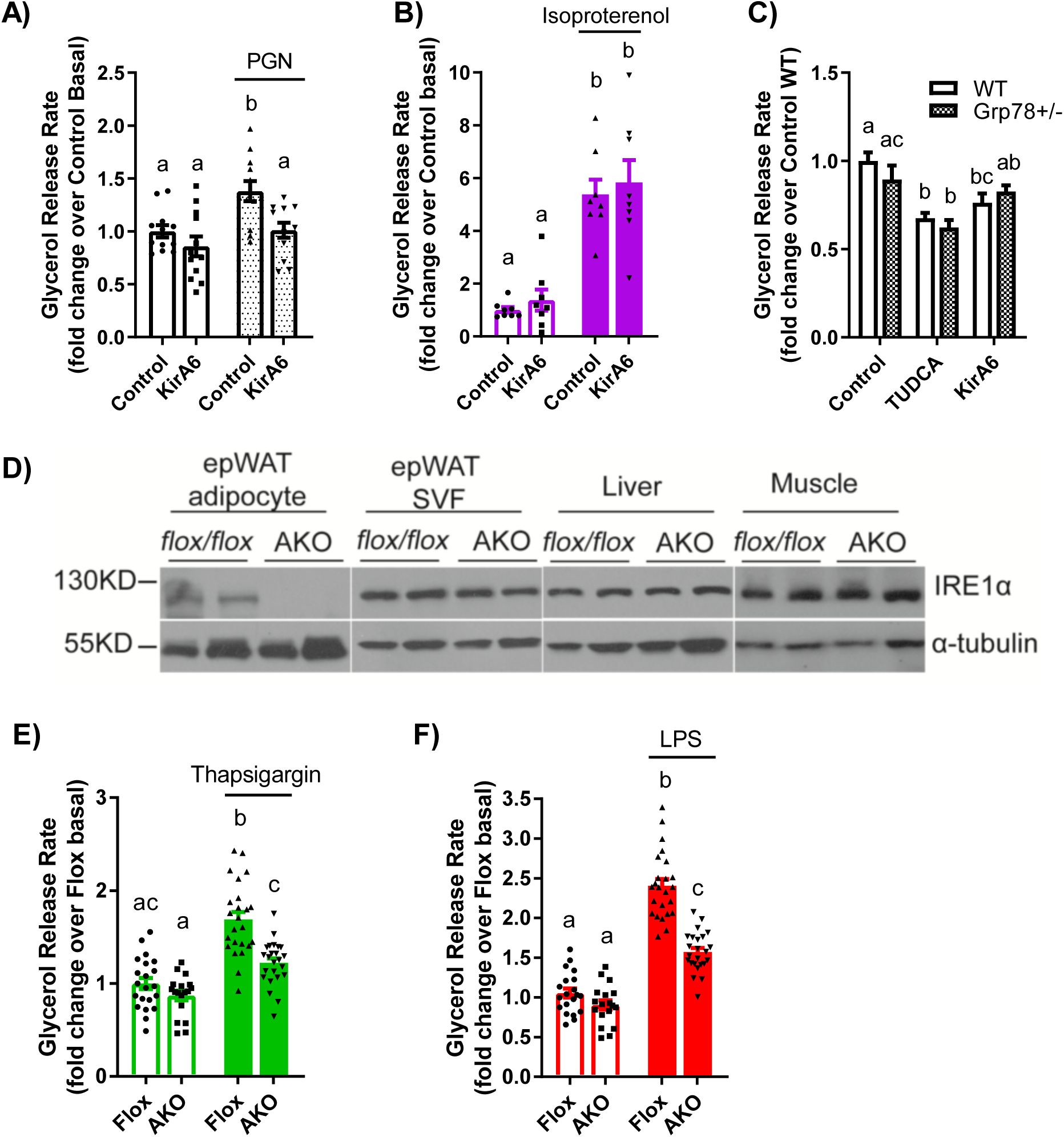
IRE1 mediates lipolysis induced by inflammatory ligands in adipose tissue. A-B) Glycerol release rates were determined in adipose tissue explants from wild type C57Bl/6J mice treated with or without PGN (10 µg/mL) or isoproterenol (2 µM) in the absence or presence of the IRE1 inhibitor KirA6 (1 µM). Glycerol release rates were expressed relative to the Control basal group (N=8-12). C) Adipose tissue explants were prepared from wild type and Grp78 heterozygous C57Bl6/J littermate mice and treated with vehicle, TUDCA (1 mg/mL), or KirA6 (1 µM) for 72 hours. Glycerol release rates were determined and expressed relative to the wild type control group (N=12). D-F) Glycerol release analysis of adipose explants from littermate control (Flox) and adipocyte-specific IRE1 knockout (AKO) mice. D) Representative immunoblots of IRE1 from adipocytes and SVF of epididymal adipose tissue, as well as from liver and muscle from Flox and AKO mice. E-F) Glycerol release rates were measured in epididymal adipose tissue explants from Flox control versus AKO mice. Explants were treated with or without 1 µM thapsigargin (E) or 500 ng/mL LPS (F) for 72 hours and glycerol release was measured and expressed relative to the Flox basal group (N=18-24 explants). Values are mean ± SEM. Statistical significance was measured as p<0.05 using two-way ANOVA. Post hoc analysis was performed using Tukey’s multiple comparisons test. Conditions with different letters (a, b) denote a statistical difference compared to all other conditions without the same letter.

### IRE1 kinase links inflammation to NF-κβ and cytokine secretion

Activation of NF-Κβ is a hallmark of inflammation and we have previously reported that bacterial cell wall components (such as PGN and other NOD1 ligands) engage NF-Κβ to stimulate adipocyte lipolysis (11). However, it is unknown if IRE1 acts upstream of NF-Κβ to potentiate further cytokine secretion to augment lipolysis. HEK293 cells, stably expressing NOD1 or TLR4 with an NF-κβ SEAP reporter, were used to assess the involvement of IRE1 in activating NF-κβ in response to inflammatory stimuli. Treatment of HEK-NOD1 cells with either PGN (10 µg/mL FK565) or TNF (10 ng/mL), or HEK-TLR4 cells with 500 ng/mL LPS, increased NF-κβ activity, whereas isoproterenol (in HEK-NOD1 cells) did not activate NF-κβ (Figure 4A-D). KirA6 completely blocked NF-κβ activation induced by PGN and partially blocked NF-κβ activation induced by TNF. KirA6 had a very small, but statistically significant, inhibitory effect on LPS-induced NF-κβ activation (Figure 4C). These data suggest that IRE1 kinase propagates inflammatory ligand-induced NF-κβ activation from NOD1 and TNFR, but biological relevance to TLR4 is not yet clear. Nevertheless, our data positions IRE1 kinase activity upstream of NF-κβ activation in responses to inflammatory triggers of lipolysis in adipocytes.

**Figure 4:**
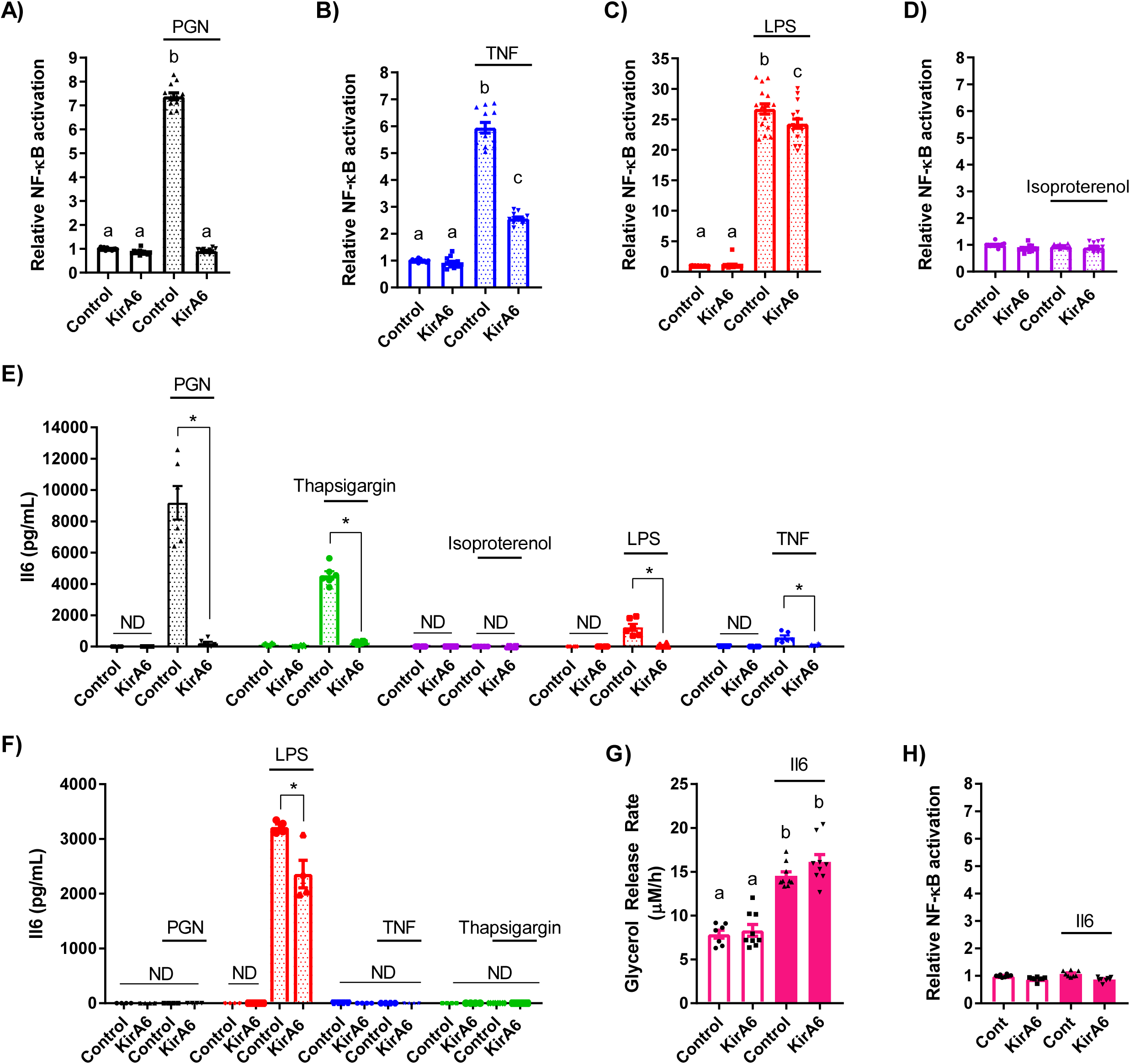
Inflammatory ligands activate NF-kB and Il6 secretion through an IRE1 pathway in 3T3-L1 adipocytes. A-D) NF-κβ activation was measured in HEK-Blue™ NOD1 (A, B, D) or TLR4 (C) cells and reported relative to the vehicle control. Cells were treated with vehicle, 10 µg PGN (A), 10 ng/mL TNF (B), 500 ng/mL LPS (C), or 2 µM isoproterenol (D), in the presence or absence of 1µM KirA6 (N=12). E) Il6 secretion from differentiated 3T3-L1 adipocytes was measured by ELISA after 48 hours of incubation in either the presence or absence of 1 µM KirA6, treated with vehicle, 10 µg/mL PGN, 1 µM thapsigargin, 2 µM isoproterenol, 500 ng/mL LPS, or 10 ng/mL TNF (N=6). F) Il6 secretion from bone-marrow derived macrophages was measured by ELISA after 48 hours of incubation with or without 1 µM KirA6 and 10 µg/mL PGN, 500 ng/mL LPS, 10 ng/mL TNF, or 1 µM thapsigargin (N=4). G) Glycerol release rates were measured in 3T3-L1 adipocytes treated with or without Il6 (20 ng/mL) in the absence or presence of 1 µM KirA6 (N=9). H) NF-κβ activation was measured via in HEK-Blue™ NOD1 cells with or without Il6 (20 ng/mL) in the absence or presence of 1 µM KirA6 (N=8). Values are mean ± SEM. Statistical significance was measured as p<0.05 using two-way ANOVA (A-D; G, H). Post hoc analysis was performed using Tukey’s multiple comparisons test. Conditions with different letters (a, b) denote a statistical difference compared to all other conditions without the same letter (A-D). Student t-test was used to determine statistical significance in panels E and F. * denotes significance p<0.05.

We next investigated if IRE1 kinase activity mediates inflammatory stimuli-induced cytokine secretion in adipocytes. Previous work has placed IRE1 upstream of NF-Κβ in regulating Il6 secretion in macrophages (20). We found that 3T3-L1 adipocytes secrete in response to the inflammatory triggers PGN, LPS, and thapsigargin. KirA6 inhibited Il6 secretion stimulated by all inflammatory stimuli (PGN, LPS, TNF, thapsigargin), whereas isoproterenol did not stimulate Il6 secretion, in 3T3-L1 adipocytes (Figure 4E). We also tested Il6 secretion in bone marrow derived macrophages (BMDMs). Only LPS stimulated Il6 secretion in BMDMs, which is consistent with previous results (22). LPS stimulated Il6 secretion was partially inhibited by KirA6 in BMDMs (Figure 4F). These data support a role for IRE1 kinase activity in mediating Il6 secretion in response to inflammatory stimuli in adipocytes.

Given the role of IRE1 in regulating Il6 secretion in response to inflammatory stimuli, we tested whether Il6 exposure induces IRE1 kinase-dependent lipolysis. Il6 increased glycerol release, but Il6-induced lipolysis was not altered by KirA6 (Figure 4G). Furthermore, Il6 did not activate NF-Κβ activity in HEK-293T cells stably expressing NOD1 and an NF-Κβ SEAP reporter (Figure 4H). Thus, our data is consistent with a model where IRE1 contributes to Il6 secretion from inflammatory triggers, but autocrine Il6 secretion does not feedback into an IRE1-dependent mechanism to stimulate lipolysis. Our data also show that not all cytokines engage IRE1 kinase to promote lipolysis, since KirA6 blocked TNF-stimulated, but not Il6-simulated, lipolysis in adipocytes.

### IRE1 kinase regulates lipolysis, independent of other tyrosine kinases

Multiple other kinases may mediate inflammation induced lipolysis. For example, the ABL kinase, c-ABL, has been directly implicated in IRE1-linked ER stress responses (23). Further, RIPK2 is a kinase required for NOD1-mediated NF-Κβ activation and immune responses and RIPK2 has also been reported to propagate ER stress responses (20, 21). We examined if these other kinase targets were involved in inflammation-induced lipolysis using the tyrosine kinase inhibitors (TKIs) ponatinib (0.1 µM) and imatinib (5 µM). Ponatinib has a high affinity for RIPK2, whereas imatinib potently inhibits c-ABL, but does not inhibit RIPK2 (24). Ponatinib completely blocked glycerol release stimulated by PGN, as expected due to its inhibition of RIPK2 (Figure 5A). However, ponatinib failed to inhibit lipolysis stimulated by any other inflammatory or adrenergic stimulus (Figure 5A, B). Imatinib inhibited thapsigargin-induced sXBP1 levels in adipocytes (Figure 5C), demonstrating that imatinib can inhibit downstream consequences of IRE1 RNase activity. However, imatinib did not affect increased lipolysis induced by any stimuli (Figure 2D). These data indicate that RIPK2 is not required for all inflammation-stimulated lipolytic programs, rather RIPK2 is only involved in the NOD1-mediated lipolytic pathway. Furthermore, RIPK2 plays no role in lipolysis stimulated by adrenergic triggers of lipolysis such as isoproterenol. More importantly, the multiple targets inhibited by imatinib, such as ABL kinases, do not discriminate inflammatory versus hormonal-induced lipolysis. In fact, we find no role for the kinase targets of ponatinib and imatinib in regulating lipolysis beyond blocking RIPK2 responses to PGN. Seminal work showed that imatinib-mediated targeting of ABL-IRE1 alleviates ER stress in pancreatic beta cells via RNase activity and a UPR (23). We showed that imatinib can inhibit the target of IRE1 RNase activity, sXBP1, in thapsigargin-treated adipocytes, but the multiple kinases inhibited by imatinib (including c-ABL) did not alter inflammation or adrenergic lipolysis.

**Figure 5:**
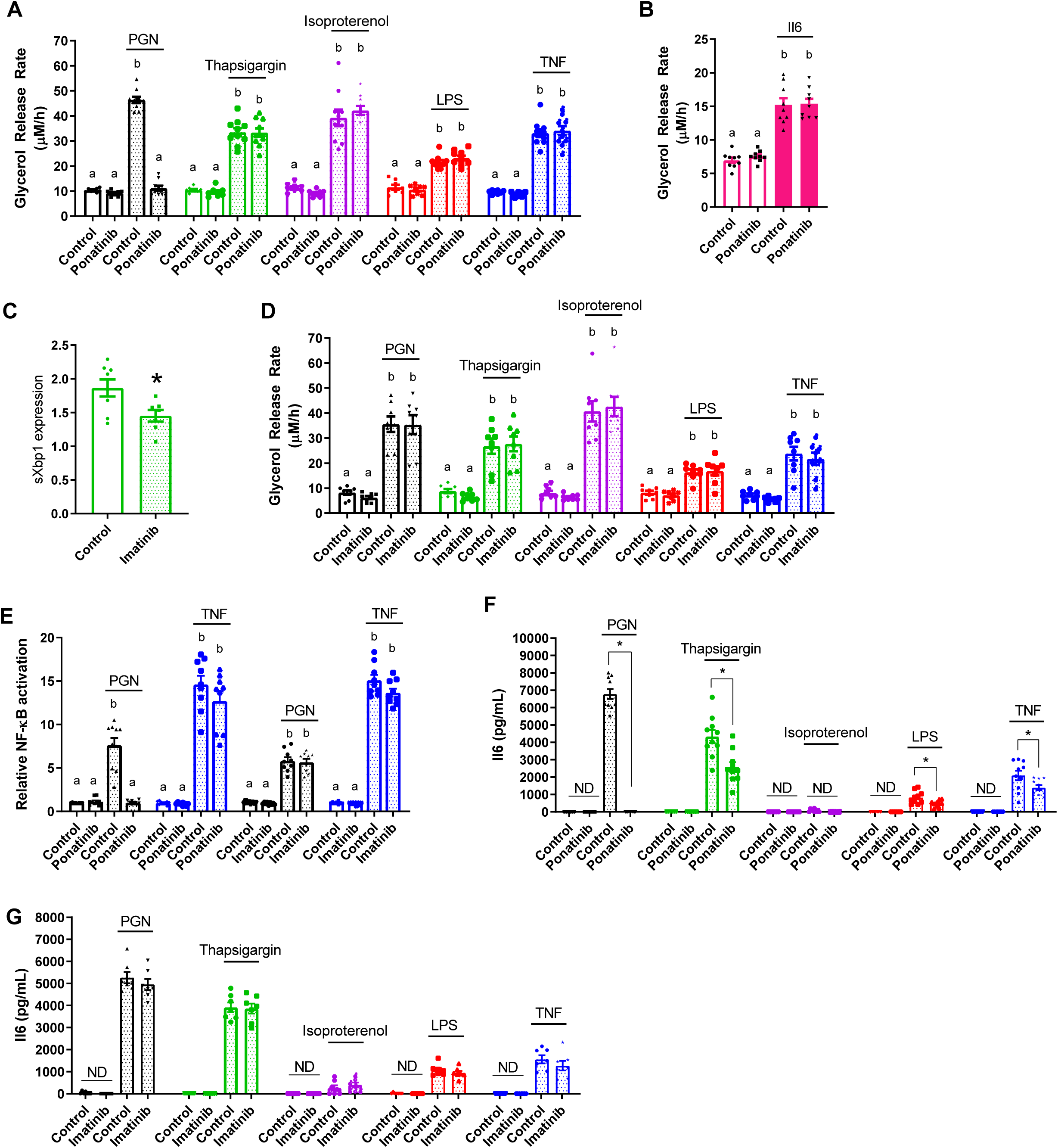
The tyrosine kinase inhibitors ponatinib and imatinib do not alter the IRE1-mediated lipolysis in 3T3-L1 adipocytes. A, B, D) Glycerol release rates were calculated in adipocytes treated with vehicle (control), 10 µg/mL PGN, 1 µM thapsigargin, 2 µM isoproterenol, 500 ng/mL LPS, 10 ng/mL TNF, or 20 ng/mL Il6, as indicated (N=8-10). Cells were treated with ponatinib (0.1 µM) or imatinib (5 µM) for 1 hour prior to ligand addition. C) Relative transcript levels of spliced XBP1 (sXBP1) in 3T3-L1 adipocytes treated with 0.25µM Thapsigargin for 3 h with or without 5 µM imatinib. E) NF-κβ activation in HEK-Blue™ NOD1 cells treated with vehicle, 10 µg/mL PGN or 10 ng/mL TNF in the presence or absence of 0.1 µM ponatinib or 5 µM imatinib (N=8-9). F-G) Il6 secretion in 3T3-L1 adipocytes after 48 hours of incubation with or without 0.1 µM ponatinib (F) or 5 µM imatinib (G) and 10 µg/mL PGN, 1 µM thapsigargin, 2 µM isoproterenol, 500 ng/mL LPS, or 10 ng/mL TNF, as indicated (N=7-10). Values are mean ± SEM. Statistical significance was measured as p<0.05 using two-way ANOVA (A, B, D, E). Post hoc analysis was performed using Tukey’s multiple comparisons test. Conditions with different letters (a, b) denote a statistical difference compared to all other conditions without the same letter. Student t-test was used to determine statistical significance in panels C, F and G. * denotes significance p<0.05.

We next examined if these TKIs affect inflammation-induced NF-κβ/cytokine responses in a different way compared to lipolysis. As expected, RIPK2 inhibition with ponatinib completely blocked PGN-stimulated NF-κβ activation, but ponatinib did not alter TNF-stimulated NF-κβ activation (Figure 5E). Imatinib did not alter NF-κβ activation induced by PGN or TNF (Figure 5E). Ponatinib completely blocked NOD1-stimulated Il6 secretion and significantly lowered Il6 secretion stimulated by thapsigargin, LPS, and TNF (Figure 5F). Imatinib did not affect Il6 secretion stimulated by any stimulus (Figure 5G). Hence, our data show that bacterial, cytokine and ER stress ligands trigger an intracellular kinase response that is different for lipolysis compared to Il6 secretion. Clearly, RIPK2 is required for PGN-NOD1-stimulated Il6 secretion and lipolysis, but our data also shows that the kinases targeted by ponatinib also contribute to the Il6 response for other inflammatory stimuli, at least at the dose of ponatinib used in adipocytes (0.1 μM). Overall, our data is consistent with a model where IRE1 kinase activity specifically regulates a lipolytic program in response to inflammatory stimuli, but other kinases are engaged to regulate immune response such as cytokine secretion.

### Inflammation via IRE1 kinase does not affect insulin signalling through PKB

Inflammation and ER stress are known to promote insulin resistance (4, 6, 15). However, it is unknown if inflammatory triggers act through IRE1 kinase to promote adipocyte insulin resistance, which would then reduce the ability of insulin to suppress lipolysis. Exposure of adipocytes to PGN or thapsigargin inhibited phosphorylation of Akt/PKB in response to insulin (Figure 6A, B). However, inhibiting IRE1 kinase activity with KirA6 during PGN or thapsigargin exposure did not alter insulin signalling. In contrast, exposure of adipocytes to isoproterenol (Figure 6C), LPS (Figure 6D) or TNF (Figure 6E) failed to inhibit insulin-stimulated phosphorylation of PKB. Thus, impaired insulin action is not a unifying factor underpinning adipocyte-autonomous lipolysis from inflammatory or adrenergic stimuli. Further, our data suggests that IRE1 kinase-dependent lipolysis in adipocytes can function independently of insulin resistance that can manifest from inflammation or ER stress.

**Figure 6:**
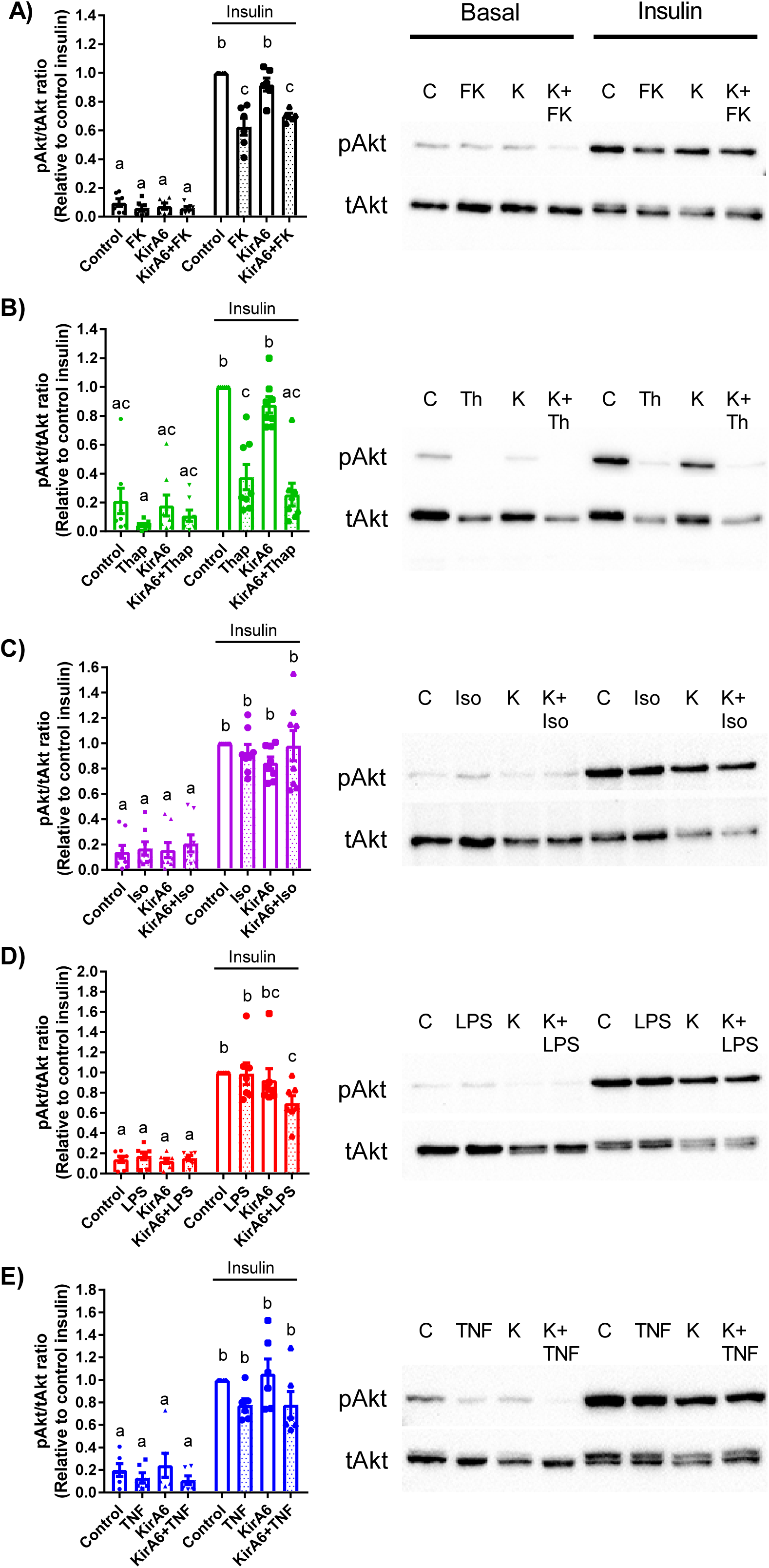
IRE1 kinase activity does not affect insulin signalling through PKB. Phosphorylation of Akt/PKB (ser473) relative to total Akt/PKB was measured by immunoblotting in 3T3-L1 adipocytes in the absence or presence of 1 µM KirA6. Cells were pre-treated with or without KirA6 for 1 hour before addition of ligand for 48 hours. Cells were treated with or without 100 nM insulin for the final 10 min after cells were exposed for 48 hours to: A) PGN (10 µg/mL), B) thapsigargin (1 µM), C) isoproterenol (2 µM), D) LPS (500 ng/mL), or E) TNF (10 ng/mL). Values are mean ± SEM (N=6-8). Statistical significance was measured as p<0.05 using two-way ANOVA. Post hoc analysis was performed using Tukey’s multiple comparisons test. Conditions with different letters (a, b) denote a statistical difference compared to all other conditions without the same letter.

## Discussion

Obesity is associated with inflammation and increased adipocyte lipolysis. Lipolysis can be increased though hormones and higher adrenergic activity or from inflammatory mediators inherent to adipose tissue during obesity. One connection between obesity-induced inflammation and hormonal control of lipolysis is insulin resistance, which manifests as impaired insulin-mediated suppression of lipolysis. Another key connection between inflammation and lipolysis showed that increased activation of ERK can stimulate Beta3 adrenergic receptor-mediated lipolysis via protein kinase A (PKA) (19). This outstanding discovery showed how a stress kinase could influence adrenergic-mediated lipolysis. Obesity is associated with increased ERK activity/phosphorylation preferentially in white adipose tissue (19). Obesity is also associated with higher insulin and increased levels of many potential triggers of inflammation within adipose tissue, including bacterial components and pro-inflammatory cytokines (3, 25). Indeed, cytokines and higher insulin can both activate ERK in adipose tissue, but it was not clear if triggers of inflammation engaged a different cellular response compared to hormones or if inflammation required insulin resistance to promote excessive lipolysis. It was also already known that ERK and PKA mediate part of the lipolytic response after adipocytes are exposed to bacterial components such as LPS or PGN (11, 12). However, it was not known if inflammatory mediators engage a specific kinase to promote a lipolytic program, which would represent a unifying factor for inflammation-induced lipolysis that is separate from hormonal triggers and adrenergic control of lipolysis.

ERK is a key regulator of HSL-mediated adipocyte lipolysis (26). However, ERK is not positioned to differentiate adrenergic and inflammatory lipolysis, rather ERK integrates cell stress and adrenergic signalling and represents a common node that promotes crosstalk between stress and PKA/cAMP-mediated adipocyte lipolysis (19). Previous work has shown that inflammatory factors require different conditions to promote lipolysis compared to adrenergic stimuli. For example, TNF requires glucose in the media to stimulate adipocyte lipolysis, whereas extracellular glucose does not alter isoproterenol-induced lipolysis (27). Intriguingly, glucose did not alter the ability of TNF to stimulate proximal signalling events in lipolysis such as ERK activation. These data highlight the need to assess lipid/glycerol release (rather than just signal activation) and warrant scrutiny of the intracellular mechanism responsible for propagation of the lipolytic response due to inflammatory mediators. We assessed if a component of ER stress discriminated inflammatory triggers of lipolysis. The kinase activity of IRE1 was required for inflammatory triggers such as bacteria or certain cytokines to promote lipolysis.

Overall, our data support a model where the kinase activity of IRE1 is the component of ER stress that mediates inflammation-induced lipolysis. Inflammatory stimuli require IRE1 kinase, but not RNase, activity to stimulate lipolysis. Adrenergic stimuli do not require IRE1 to promote lipolysis. Specific receptors for each inflammatory trigger converge on IRE1 to promote lipolysis. NOD1 activation requires RIPK2 to promote IRE1-mediated lipolysis and Il6 secretion. Not all cytokines take the same path to promote lipolysis, since TNF engages IRE1 and NF-κβ to increase lipolysis, but Il6 can circumvent these requirements.

Our results implicate IRE1 kinase activity in NF-κB and immune responses triggered by bacteria and certain cytokines, such as TNF. PKA can influence NF-κB and we already knew that ERK, PKA and NF-κB conspired to promote adipocyte lipolysis (11, 28). Our data show that IRE1 kinase activity is required for certain bacterial and cytokine triggers of immune responses to activate NF-κB. IRE1 kinase control of NF-κB is positioned to propagate lipolysis, but also a plethora of immune and stress responses related to NF-κB activation. It was important to define the role of IRE1-linked ER stress in regulating NF-κB and cytokines secretion because it was reported that the bacterial cell wall muropeptide sensors NOD1 and NOD2 are mediators of ER stress-induced cytokine secretion in macrophages (20, 21) and in the liver (29). It was shown that thapsgargin-induced ER stress promoted macrophage secretion of Il6, which was dependent upon IRE1, NOD1/2, and RIPK2 (20, 21). Our data show that the lipolytic and inflammatory pathways share dependence on IRE1, but do not share dependence on RIPK2. Our data show that RIPK2 does not function downstream of ER stress to promote Il6 secretion in adipocytes; rather, RIPK2 only mediates immune and lipolytic responses engaged by NOD1 or NOD2. Our results using a TKI that targets RIPK2 (i.e. ponatinib) reinforce the concept that RIPK2 dictates immune and lipolysis responses to PGN acting on NOD proteins. We also used the TKI imatinib to test the role of ABL kinases in ER stress-related lipolysis and immunity. This was important given the seminal work showing that inhibiting c-ABL with imatinib lowers IRE1-mediated RNase activity during the UPR (23). Our data show no role for the UPR or RNase activity of IRE1 in the regulation of lipolysis triggered by inflammatory mediators. We found no correlation between changes in insulin signalling and lipolysis and blocking IRE1 kinase activity had no effect on insulin signalling. Our data supports a model where bacterial and cytokine inflammatory stimuli promote lipolysis, NF-κB activation, and Il6 secretion through the kinase activity of IRE1. The distinction between the hormonal/adrenergic and inflammatory triggers of lipolysis is defined by IRE1 kinase.

## Materials and Methods

### Materials

Heptanoyl-γ-D-glutamyl-L-meso-diamino-pimelyl-D-alanine (FK565) was obtained from Fujisawa Pharmaceuticals (Osaka, Japan). Fatty acid-free bovine serum albumin (BSA; #A8806), thapsigargin (#T9033), isoproterenol (# I6504), Atglistatin (#SML1075), 3-isobytyl-1-methylanthine (#I5879), dexamethasone (#D4902), rosiglitazone (#R2408), insulin solution from bovine pancreas (insulin; #I0516), sodium tauroursodeoxycholate (TUDCA; #T0266), sodium phenylbutyrate (4-PBA; SML0309) and 8-Br-cAMP (#B7880) were from Sigma-Aldrich (St. Louis, MO). Ponatinib (#A10080) and imatinib mesylate (imatinib; #A10468) were obtained from AdooQ Bioscience (Irvine, CA). Tumour necrosis factor (TNF; #575202) and Il6 (#575702) were from BioLegend (San Diego, CA). Dulbecco’s Modification of Eagle’s Medium (DMEM), Dulbecco’s phosphate-buffered saline (PBS), fetal bovine serum (FBS), and penicillin and streptomycin antibiotic solution (pen/strep) were obtained from Wisent (St Bruno, CA). GlutaMax, Restore Stripping Buffer, bicinchonic acid (BCA) assay kit, SuperScript III (#18080085), and Amplitaq Gold (#4317742) were obtained from Thermo Fisher Scientific (Waltham, MA). Kinase inhibiting RNase Attenuator 6 (KirA6; #5322810001) and IRE1 inhibitor III 4µ8C (#412512) were obtained from Calbiochem. IRE1 RNase activity inhibitor KirA8 (#HY-114368) was obtained from MedChemExpress. Lipopolysaccharide (LPS; #tlrl-ppglps), Normocin™ (#ant-nr-1), Blasticidin™ (#ant-bl-1), Zeocin™ (#ant-zn-1), HEK-Blue™ Detection media (#hb-det3), HEK-Blue™ mNOD1 (#hkb-mnod1) cells and HEK-Blue™ mTLR4 (#hkb-mtlr4) cells were obtained from InvivoGen (San Diego, CA). Antibodies for phospho-PKB (#4058) and total PKB (#9272) were obtained from Cell Signaling (Danvers, MA). Antibodies for phospho-S724-IRE1 (#ab48187) and total IRE1 (#ab37073) were from Abcam (Cambridge, MA).

### Differentiation and treatment of 3T3-L1 adipocytes

Murine 3T3-L1 pre-adipocytes were cultured in DMEM supplemented with 10% FBS, 1% GlutaMax, and 1% p/s (growth media) and maintained in an incubator at 37 °C and 5% CO2. Cells were seeded into 24-well plates in growth media and switched into differentiation media (growth media containing 0.5 mM 3-isobutyl-1-methylxanthine, 0.25 µM dexamethasone, 2 µM rosiglitazone, and 1 µg/mL insulin) 24 hours post-confluence. After 48 hours (Day 0-2), cells were incubated in growth media containing 1 µg/mL insulin for 96 hours (day 3 to 6), with the media changed on Day 5. On Day 6 cells were placed in growth media and media was changed every 48 hours until full differentiation was achieved. Experiments were performed between days 8 to 10, depending on differentiation efficiency.

At time of treatment, 3T3-L1 adipocytes were washed once with PBS and treated with or without indicated inhibitors for 1 hour in serum-free DMEM containing 0.5% fatty acid-free BSA and 1% p/s. Cells were treated with the general ER-stress inhibitors TUDCA or 4-PBA (varying concentrations), the Adipose Triglyceride Lipase (ATGL) inhibitor Atglistatin (5 or 50 µM), the IRE1 inhibitor KirA6 (1 µM, unless otherwise stated), the IRE1 RNase inhibitor KirA8 (10 µM, unless otherwise stated) or 4µ8C (30 or 60 µM) or the RIPK2 tyrosine-kinase inhibitor ponatinib (0.1 µM), or the tyrosine-kinase inhibitor imatinib (5 µM). After this 1 hour incubation one of the following ligands was added to the cells: PGN = 10 µg/mL FK565 (NOD1 activation), 500 ng/mL LPS (TLR4 activation), 10 ng/mL TNF, 20 ng/mL Il6, 1 µM thapsigargin (ER stress activation), 2 µM isoproterenol (β adrenergic activation), 0.5 mM 8-Br-cAMP, or a mock-treatment.

### Glycerol Release Assay

Following treatments of 3T3-L1 adipocytes, supernatant samples were collected at 0, 24, and 48 hours of ligand treatment (unless otherwise stated) and stored at -20 °C. Lipolysis was assessed *in vitro* by measuring the release of glycerol into the supernatant media over time (0, 24, 48 h) and calculating the glycerol release rate (µM/h). Glycerol concentration was measured as per the manufactures protocol using the free glycerol determination kit (Sigma-Aldrich, #FG0100).

### Detection of IRE1 and PKB by Western blotting

For determination of IRE1 phosphorylation, 3T3-L1 adipocytes were treated with or without the KirA6 inhibitor for 1 hour and then the ligands FK565, thapsigargin, or isoproterenol were added for 3 hours. Cells were lysed in lysis buffer containing 250mM NaCl, 50mM NaF, 5mM EDTA, 10mM Na_4_P_2_O_7_, 1mM Na_3_VO_4_, 1% TX-100, 1 complete tablet of protease inhibitor, 50mM Tris-HCl and pH 7.4. Protein concentration was measured using a BCA assay kit (Sigma Aldrich) and spectrometry at absorbance 562nm using a Synergy H4 Hybrid Reader (Biotek). Phospho-IRE1 was measured by Western blot using 20 µg protein loading per sample. Sample PVDF membranes were incubated in anti-phospho-S724-IRE1 (1:1000) antibody at 4°C overnight and in goat anti-rabbit IgG horseradish peroxidase conjugated secondary (1:5000) antibody for 1 hour before detection using ECL detection (BioRad) and ChemiDoc Imaging (BioRad). PVDF membranes were stripped and re-probed for total IRE1 (1:1000). Phospho-IRE1 signal was quantified as the ratio between phospho/total IRE1 densitometry using ImageLab software (BioRad). Relative values were normalized to the control basal band of each membrane.

For determining phospho-PKB, 3T3-L1 adipocytes were treated with or without the KirA6 inhibitor for 1 hour and then the ligands FK565, thapsigargin, isoproterenol, LPS, or TNF were added for 48 hours. Cells were washed 1x with PBS and then treated with or without 100 nM insulin for 10 min before lysis. Protein concentration was detected as above, and phospho-S473-PKB (1:1000) was measured by Western blotting using 20 µg protein loading per sample. PVDF membranes were stripped and re-probed for total PKB (1:1000). Phospho-PKB signal was quantified as the ratio between phospho/total PKB densitometry using ImageLab software and relative values were normalized to the control insulin band of each membrane.

### Quantitative PCR (qPCR) detection of transcripts

For determining mRNA levels of select transcripts, 3T3-L1 adipocytes were treated with or without the KirA6 inhibitor for 1 hour and then the ligands FK565, thapsigargin, or isoproterenol were added for 3 hours. Cells were washed 1x in PBS before being suspended in Trizol. RNA was isolated and cDNA synthesis. Amplitaq Gold (Thermo Fisher Scientific) was used for qPCR using Taqman primers to BiP (Mm00517691), CHOP (Mm01135937), and sXbp1 (Mm03464496) in a Rotor-Gene-Q real-time PCR cycler (Qiagen). The gene Rplp0 (Mm01974474) was used as reference to calculate delta Ct values. Gene expression was expressed as fold change relative to basal control samples.

### Il6 ELISA

Media samples collected at 48 hours of ligand treatment were analyzed for Il6 secretion. Il6 concentration was determined by using the Mouse Il6 DuoSet® ELISA (R&D Systems, #DY406-05), as per the manufactures protocol with the following modifications: the capture antibody was utilized at a working concentration of 1 µg/mL and the detection antibody was 75 ng/mL.

### Determination of NF-κB Activation

HEK-Blue™ NOD1 and HEK-Blue™ TLR4 cells stably express a secreted alkaline phosphatase (SEAP) reporter of NF-ΚB activity. HEK-Blue™ NOD1 cells were grown in media containing DMEM, 10% FBS, 1% GlutaMax, 1% p/s, 100 µg/mL Normocin™, 30 µg/mL Blasticidin™ and 10 µg/mL Zeocin™ (HEK-NOD1 media) and incubated at 37 °C and 5% CO_2_. HEK-Blue™ TLR4 cells were grown in media containing DMEM, 10% FBS, 1% GlutaMax, 1% p/s, 100 µg/mL Normocin™, and 1x HEK-Blue™ selection (HEK-TLR4 media). Cells were seeded at a density of 30,000 cells/well into a 96-well plate and incubated for 24 hours. Post-incubation, cells were exposed to HEK-Blue™ detection media supplemented with 1 µM KirA6, 0.1 µM ponatinib or 5 µM imatinib, as appropriate. Cultures were allowed to equilibrate for 1 hour before being treated with 10 µg/mL FK565, 500 ng/mL LPS, 10 ng/mL TNF, 20 ng/mL Il6, 1 µM thapsigargin, 2 µM isoproterenol, or a mock-treatment. After 24 hours, SEAP activity was measured by absorbance at 630 nm using the Synergy H4, Hybrid Reader (BioTek).

### Adipose tissue explants and generation of adipose specific IRE1 knock out mice

Male C57Bl6/J mice aged 10-12 weeks were used for adipose tissue explant experiments. These animal experiments were approved by McMaster University Animal Ethics Review Board. For experiments using Grp78^+/-^ mice, the WT controls were paired littermates. Gonadal fat pads were excised, cut into pieces ∼3-4 mg in size, and placed in DMEM containing 10% FBS and 1% pen/strep for 1 hour at 37°C and 5% CO_2_. Explanted tissues were then transferred to a second dish of DMEM containing 1% BSA and 1% pen/strep for 30 minutes. Next, 3-4 pieces of adipose tissue explant were placed per well into 24-well plates and treated with or without inhibitor (KirA6 or TUDCA) for 1 hour. After 1 hour, ligand was added to stimulate lipolysis and supernatant was collected at 0, 24, 48, and 72 hours (inflammatory ligands) or at 0, 10, 30, 60 min (isoproterenol) for assessment of glycerol release. Explanted tissue was collected, dried, and weighed for normalization of the glycerol release data.

Adipocyte-specific IRE1 knockout (AKO) mice on the C57BL/6J background were generated by crossing *Adiponectin-Cre* mice (30) with *Ern1*^*flox/flox*^ mice (31). Epididymal fat pads were isolated from 8 week old male AKO and *Ern1*^*flox/flox*^ control mice, cut into 3-4 mg pieces, and then placed in DMEM (high glucose) containing 10% FBS and 1% pen/strep for 1 hour at 37°C and 5% CO_2_. Fat explants were then transferred and incubated in DMEM containing 1% BSA (Sigma-Aldrich, A8806) and 1% pen/strep for 30 minutes before seeding into a 24-well plate with 3-4 pieces of explants per well in 1 ml DMEM (1% BSA and 1% pen/strep). After incubation for 1 hour, Thapsigargin or LPS were added and the supernatants were collected at the indicated time for assessment of glycerol release. Explants were collected, dried, and weighed for normalization. Supernatant glycerol concentration was measured using the free glycerol determination kit (Sigma-Aldrich, F6428) and glycerol standard (Sigma-Aldrich, G7793). These animal experiments were performed according to protocols approved by the Animal Care and Use Committee at the College of Life Sciences, Wuhan University.

### Bone Marrow-Derived Macrophage (BMDM) isolation and Il6 secretion

Bone marrow from the tibia and femur of C57BL6/J mice was harvested by flushing bones with DMEM and collecting the resultant cell suspension. Cells were centrifuged at 3500g at 4 °C for 5 minutes and pelleted cells were washed in DMEM twice before plating in a 175cm sq. flask for 3h in Macrophage Culture Media (MCM; DMEM + 10%FBS + 1% p/s + 15% L929-conditioned media). After 3 hours, cells were gently washed, and adherent cells were discarded. Non-adherent cells were plated in 24-well plates at 2.5×10^5^ cells per well in MCM. L929-conditioned media was replenished on Day 3-4 (15% of total well volume) and cells were used for experiments between Day 7-10. Cells were treated with or without inhibitor (KirA6 or ponatinib) for 1 hour before addition of inflammatory ligand (FK565, LPS, TNF, or thapsigargin). Media was collected after 48 hours and Il6 secretion was tested by ELISA.

### Statistical Analysis

Data are represented as the mean ± standard error of the mean (SEM). Comparisons were made using an unpaired, two-tailed Student’s t-test or two-way ANOVA and Tukey’s post hoc analysis, as appropriate using GraphPad Prism 7 software. Differences between groups were considered statistically significant when the P-value was less than 0.05.

## Acknowledgments

This work was supported by grants to J.D.S. from the Canadian Institutes of Health Research (FDN-154295) and to Y.L. from the National Natural Science Foundation of China (No. 31690102). A.K.T. was supported by ICMR International Fellowship from the Indian Council of Medical Research, Govt. of India. J.D.S. holds a Canada Research Chair in Metabolic Inflammation.

## Abbreviations

8-Br-cAMP: 8-Bromoadenosine 3′,5′-cyclic monophosphate
4-PBA: 4-Phenylbutyric acid
ABL: Abelson murine leukemia viral oncogene homolog
AKO: adipose knock out
ATF6: activating transcription factor 6
ATGL: adipose triglyceride lipase
BMDM: bone marrow derived macrophage
CHOP: C/EBP homologous protein
ER: endoplasmic reticulum
ERK: Extracellular Signal-Regulated Kinase
FK565: Heptanoyl-γ-D-glutamyl-L-meso-diamino-pimelyl-D-alanine
Grp78/BiP: binding immunoglobulin protein or heat shock 70kDa protein 5
HEK: human embryonic kidney
HSL: hormone sensitive lipase; Il(6) = interleukin
IRE1: inositol-requiring enzyme 1
KirA6: IRE1α Kinase Inhibiting RNase Attenuator 6
NF-ΚB: nuclear factor kappa-light-chain-enhancer of activated B cells
LPS: lipopolysaccharide
NOD: Nucleotide-binding oligomerization domain-containing protein
PERK: protein kinase RNA-like ER kinase
PGN: peptidoglycan
PKA: protein kinase A
PKB/Akt: protein kinase B
RIPK2: Receptor Interacting Serine/Threonine Kinase 2
SEAP: secreted alkaline phosphatase
SVF: stromal vascular fraction
TLR4: toll-like receptor 4
TKI: tyrosine kinase inhibitor
TNF: tumor necrosis factor
TUDCA: Tauroursodeoxycholic acid
UPR: unfolded protein response
WAT: white adipose tissue
WT: wild type
XBP1: X-Box Binding Protein 1

